# The ability to sense the environment is heterogeneously distributed in cell populations

**DOI:** 10.1101/2023.03.07.531554

**Authors:** Andrew Goetz, Hoda Akl, Purushottam Dixit

## Abstract

Channel capacity of signaling networks quantifies their fidelity in sensing extracellular inputs. Low estimates of channel capacities for several mammalian signaling networks suggest that cells can barely detect the presence/absence of environmental signals. However, given the extensive heterogeneity and temporal stability of cell state variables, we hypothesize that the sensing ability itself may depend on the state of the cells. In this work, we present an information theoretic framework to quantify the distribution of sensing abilities from single cell data. Using data on two mammalian pathways, we show that sensing abilities are widely distributed in the population and most cells achieve better resolution of inputs compared to an “*average cell*”. We verify these predictions using live cell imaging data on the IGFR/FoxO pathway. Importantly, we identify cell state variables that correlate with cells’ sensing abilities. This information theoretic framework will significantly improve our understanding of how cells sense in their environment.

## Introduction

In cell populations, there is a significant overlap in responses to environmental stimuli of differing strengths. This raises a fundamental question^1^: do signaling networks in cells relay accurate information about their environment to take appropriate action? And if not, where along the signal transduction pathway is the information lost^2, 3^? Mutual information (MI) quantifies the information content in an intracellular output about extracellular inputs. For an input *u* (e.g., concentration of a ligand) and an output *x* (e.g., intracellular species concentration^4-6^ or a cellular phenotype^7, 8^), the MI is defined as^9^

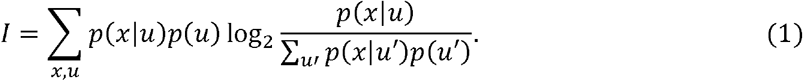

Experimental single cell methods such as flow cytometry^10^, immunofluorescence^10^, mass spectrometry^11^, or live cell imaging^12^ allow us to estimate response histograms *p*(*x*|*u*) across several inputs. Using these distributions, we can estimate the maximum of the MI (the channel capacity, CC) by optimizing Eq. 1 over input distributions *p*(u).The CC quantifies fidelity of signal transduction^1, 2^. For example, a CC of 1 bit implies that that the cells can at best distinguish between two input levels (e.g. presence versus absence), with higher CCs indicating that cells can resolve multiple input states. Importantly, CC can be used to identify bottlenecks in signaling^2, 13^.

The CC has been estimated for input-output relationships in several mammalian signaling networks^4-6, 14-16^. When the output is defined as levels of a single protein a fixed time, the CC was found to be surprisingly low, ∼1 - 1.5 bits. These estimates have been improved by considering multidimensional outputs^5^ or time varying inputs^17^. While these modifications led to somewhat higher CC estimates, the overall conclusion that cells know little about their environment remains well established. In contrast, significantly higher CC estimates have been found when the output at the level of cell population averages are considered^6^, suggesting that the only way to overcome low sensing fidelity is population average response.

These previous calculations estimated one channel capacity for all cells in a population, implicitly assuming that individual cells have similar sensing abilities. However, we now know that cells exhibit extensive heterogeneity in cell state variables^18, 19^ such as abundances of key signaling proteins^20, 21^, mRNA abundances^22^, and chromatin conformation^23^ and accessibility^24^. Notably, the time scale of fluctuations in these variables can be significantly slower than relevant signaling time scales^25, 26^, sometimes extending across multiple generations^27^. Heterogeneity in cell state variables leads to a heterogeneity in response to extracellular cues, including chemotherapeutic drugs^25, 26, 28^, mitogens^29^, hormones^30^, chemotactic signals^7, 27, 31, 32^, and other electrical and chemical stimuli^33, 34^. Therefore, we hypothesize that that the ability to sense the environment varies from cell-to-cell in populations, in a cell state dependent manner.

There is no conceptual framework to estimate the variation in sensing abilities in cell populations and its dependence on cell state variables. To that end, we introduce an information theoretic quantity CeeMI: **Ce**ll stat**e** dependent **M**utual Information. We show that using typically collected single cell data and computational models of signaling networks, we can estimate the distribution *p*_CeeMI_ (*I*)of single cell signaling fidelities (single cell mutual information values). We also show that we can identify cell state variables that make some cells better and others worse at sensing their environment.

Using an illustrative example, we show that in heterogeneous cell populations, estimates of mutual information that average over cell states can be significantly lower than the mutual information of signaling networks in typical cells. Next, using previously collected experimental data, we estimate *p*_CeeMI_(*I*) for two important mammalian signaling pathways^35^; the Epidermal growth factor (EGF)/EGF receptor pathway and the Insulin like growth factor (IGF)/Forkhead Box protein O (FoxO) pathway, we show that while the cell state agnostic CC estimates for both pathways are ∼ 1 bit, most individual cells are predicted to be significantly better at resolving different inputs. Using live cell imaging data for the IGF/FoxO pathway, we show that our estimate of variability in sensing abilities matches closely with experimental estimates. Importantly, for this pathway, we also verify our prediction that specific cell state variables dictate cells’ sensing abilities. Finally, using a simple receptor/ligand model, we show how the time scales cell state dynamics affects cells’ individuality and therefore sensing abilities. We believe that CeeMI will be an important tool to explore variability in cellular sensing abilities and in identifying what makes some cells better and others worse in sensing their environment.

## Results

### Conditional mutual information models single cells as cell state dependent channels

Consider a cell population where cells are characterized by state variables ***θ***. These include abundances of signaling proteins and enzymes, epigenetic states, etc. We assume that cell states are temporally stable, that is, ***θ*** remains constant over a time scale that is longer than typical fluctuations in environmental inputs. Later, using a toy simple model, we explicitly study the effect of cell state dynamics on cells’ ability to sense their environment.

We assume that cell state variables are distributed in the population according to a distribution *p*(***θ***). If *x* denotes an output of choice (e.g., phosphorylation levels of one or more protein(s) at one or more time point(s)) and *u* denotes the input (e.g., ligand concentration), the experimentally measured response distribution *p*(*x*|*u*) can be decomposed as:

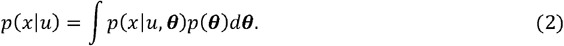

where *p*(*x*|*u, **θ***)is the distribution of the output *x* conditioned on the input *u* and cell state variables ***θ***. We note that in most cases, the same cell cannot be probed multiple times. Consequently, *p*(*x*|*u, **θ***)may not be experimentally accessible. However, it is conceptually well defined and mathematically accessible when interactions amongst molecules are specified^36^.

Using *p*(*x*|*u, **θ***), we can define the cell state dependent mutual information *I*(***θ***) for a fixed input distribution *p* (*u*) analogously to Eq. 1:

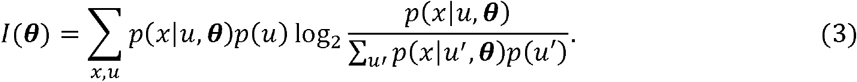

*I*(***θ***) quantifies individual cells’ ability to sense their environment as a function of the cell state parameters ***θ***. The distribution *p*_CeeMI_ (*I*) of single cell sensing abilities is

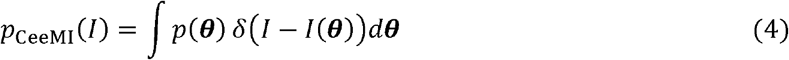

where *δ*(·)is the Dirac delta function. We can also compute the joint distribution between the single cell mutual information and any cell state variable of interest χ(e.g., abundance of cell surface receptors):

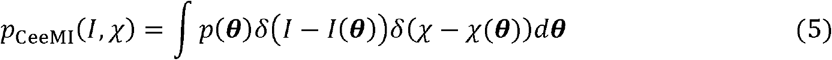

where χ(***θ***) is the value of the biochemical parameter when cell state variables are fixed at ***θ***. The distributions in Eq. 5 quantify the interdependency between a cell’s signaling fidelity *I*(***θ***) and cell specific biochemical parameters χ (***θ***). As we will see below, the distributions in Eq. 4 and Eq. 5 can be experimentally verified when appropriate measurements are available. Finally, we define the population average of the cell state dependent mutual information:

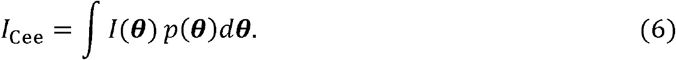

In information theory, *I*_Cee_ is known as the conditional mutual information^9^ between the input *u*and the output *x* conditioned on ***θ***. If cell state variables remain constant and if distribution over cell states is independent of the input distribution, it can be shown that *I*_Cee_ ≥ *I* (SI Section 1). It then follows that at least some cells in the population have better sensing abilities compared to the cell state agnostic mutual information (Eq. 1).

Since *I*_Cee_ depends on the input distribution *p*(*u*),we can find an optimal input distribution *p*(*u*)that maximizes *I*_Cee_ (SI Section 1). Going forward, unless an input distribution is specified, the distributions *p*_CeeMI_ (*I*) and *p*_CeeMI_ (*I, χ*)are discussed in the context of this optimal input distribution.

### Maximum entropy inference can estimate *p*_CeeMI_ (*I*)

Estimation of *p*_CeeMI_ (*I*)requires decomposing the experimentally observed response *p*(*x*|*u*) into cell specific output distributions *p*(*x*|*u, **θ***)and the distribution of cell state variables *p*(***θ***) (Eq. 3 and 4). This problem is difficult to solve given that neither *p*(*x*|*u, **θ***)nor *p*(***θ***) are accessible in typical experiments. However, for many signaling networks, stochastic biochemical models can be constructed to obtain the cell specific output distribution *p*(*x*|*u, **θ***). Here, ***θ*** comprise protein concentrations and effective rates of enzymatic reactions and serve as a proxy for cell state variables. Given the experimentally measured cell state averaged response *p*(*x*|*u*)and the model predicted cell specific output distribution *p*(*x*|*u, **θ***), we need a computational method to infer *p*(***θ***) (see Eq. 2). The problem of inference of parameter heterogeneity is a non-trivial inverse problem^37^. While there are several proposed methods to solve this problem (reviewed in^38^), most cannot efficiently infer *p*(***θ***) for signaling networks with even a moderate (n∼ 10) number of parameters. Here, we use our previously develop maximum entropy-based approach to infer *p*(***θ***)^37^. This way, we can estimate *p*_CeeMI_ (*I*) using experimentally obtained cell state agnostic response *p*(*x*|*u*) and a stochastic biochemical model *p*(*x*|*u, **θ***) of the underlying signaling network.

### An “average cell” can discern much less than a typical cell about the environment

To illustrate the effect of heterogeneity of cell state variables on the cell state agnostic estimate of mutual information (which we call *I*_CSA_ from now onwards), we consider a simple stochastic biochemical network of a receptor-ligand system. Details of the model and the calculations presented below can be found in SI Section 2. Briefly, extracellular ligand L binds to cell surface receptors R. Steady state levels of the ligand bound receptor *B* is considered the output. The signaling network is parametrized by several cell state variables ***θ*** such as receptor levels, rates of binding/unbinding, etc. For simplicity, we assume that only one variable, steady state receptor level *R*_0_ in the absence of the ligand, varies in the population. Calculations with variability in other parameters are presented in SI Section 2.

In this population, cells’ response *p*(*B*|*L,R*0) is distributed as a Poisson distribution whose mean is proportional to the cell state variable R_0_ (SI Section 2). That is, when all other parameters are fixed, a higher R_0_ leads to lower noise (coefficient of variation). To calculate cell state dependent mutual information (Eq. 3), we assume that *p*(*L*) is a gamma distribution. As expected, *I(*R_0_)(Eq. 3) between the output *B* and the input *L* increases monotonically with *R*_0_ (inset in Fig. 2A). Moreover, given that R_0_ varies in the population (also assumed to be a gamma distribution), the sensing ability varies in the population as well (Fig. 2A). Notably, the average *I*_Cee_ of *I(*R_0_) remains relatively robust to variation in. At the same time, the traditional estimate, which is the mutual information between the input and the cell state agnostic population response (response of the “average” cell, Eq. 1 and Eq. 2), decreases as the population heterogeneity in increases. Importantly, is significantly lower than the sensing ability of most cells in the population (Fig. 2A). This is because the overlap in the population response distributions is significantly larger than that in single cell response distributions (Fig. 2B). This simple example illustrates that the traditional mutual information estimates may severely underestimate cells’ ability to resolve inputs, especially when cell state variables are heterogeneously distributed. Moreover, it is crucial to explicitly account for heterogeneity in cell state variables when estimating fidelity of cellular communication channels.

**Figure 1.**
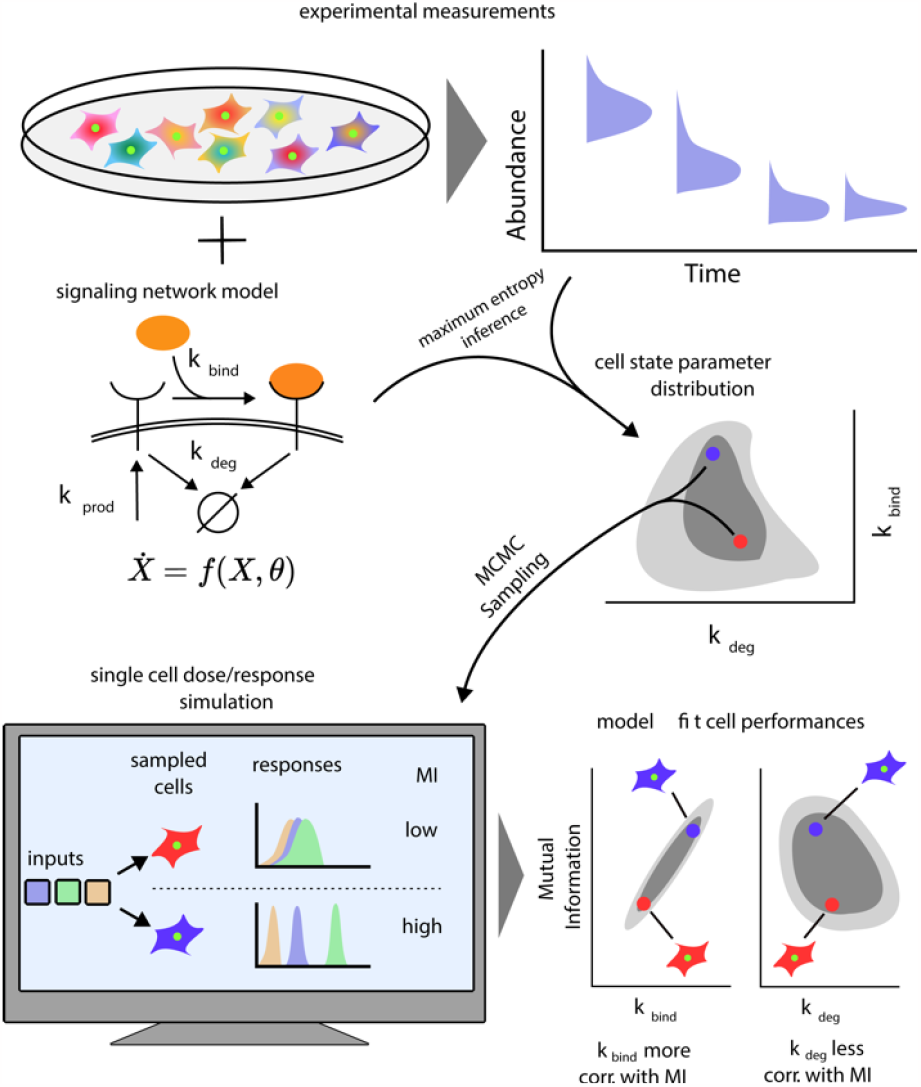
A schematic of our computational approach. (top) single cell data across different input conditions and time points are integrated with a stochastic model of a signaling network using a previous developed maximum entropy approach leading to a distribution over signaling network parameters *p*(***θ***) (middle). (bottom) In silico cells are generated using the inferred parameter distribution and cell-state specific mutual information *I*(***θ***) and population distribution of cell performances *p*_CeeMl_(*I*) is estimated. The model also evaluates the correlation between cells’ performance and biochemical parameters.

**Figure 2.**
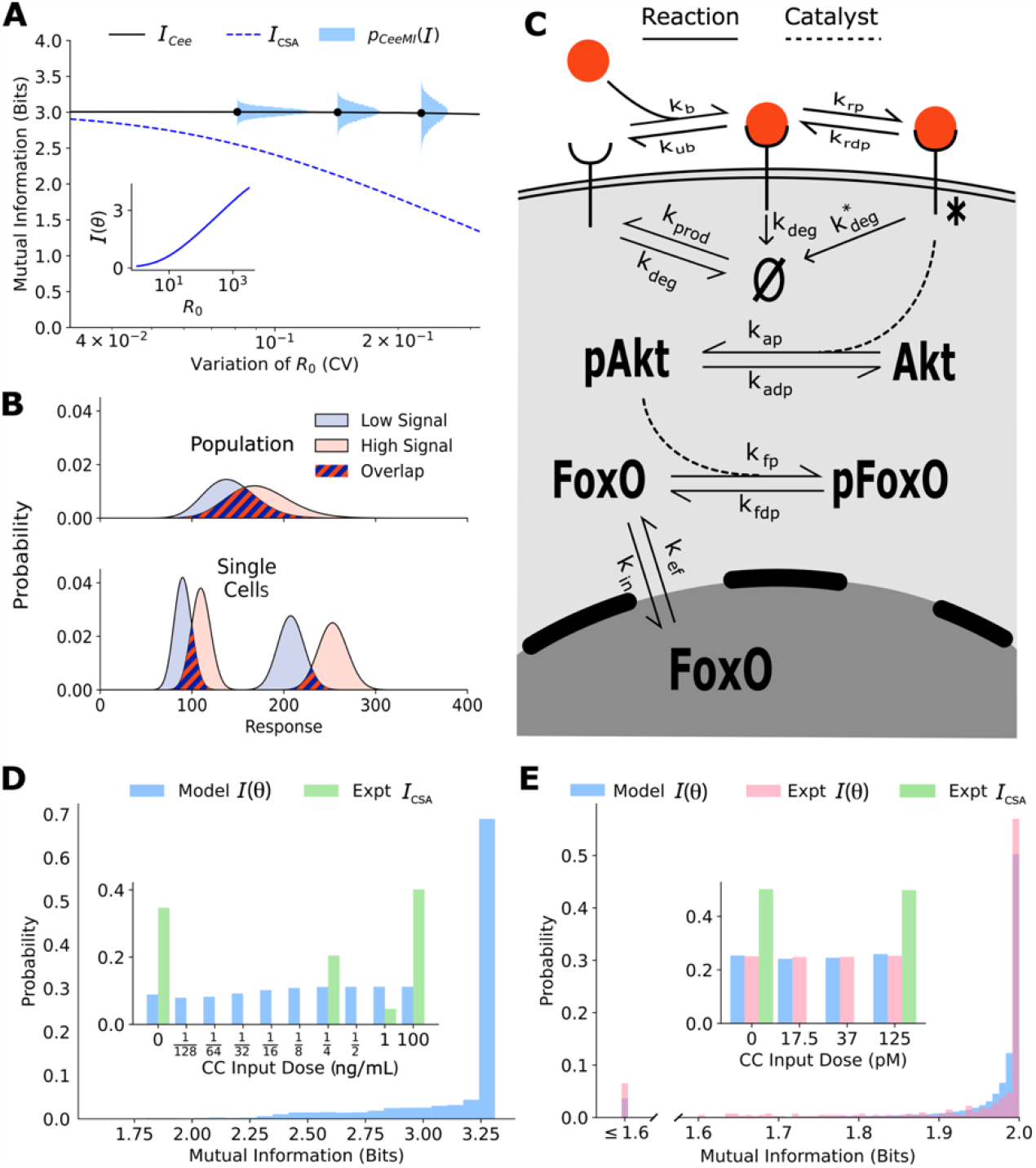
**A**. The distribution of single cell sensing abilities (horizontal blue histograms) and its averag plotted as a function of the coefficient of variation of the distribution of one cell state variable, the cell surface receptor number The dashed blue lines show the traditional cell state averaged mutual information (Eq. 1). The inset shows the dependence between cell state specific mutual information and cell state variable The input distribution is assumed to be a gamma distribution. **B**. A schematic showing the effect of heterogeneity in cell states on population level response. Even when individual cells have little overlap in their responses to extracellular signal (bottom), the population level responses could have significant overlap (top), leading to a low mutual information between cell state averaged response and the input. **C**. A combined schematic of the two growth factor pathways. Extracellular growth factor ligand (red circle) binds to cell surface receptors which are shuttled to and from the plasma membrane continuously. Ligand bound receptors are phosphorylated and activate Akt. Phosphorylated Akt leads to phosphorylation of FoxO which bars it from entering the nucleus. EGF/EGFR model is limited to the reactions on the plasma membrane. The corresponding cell state variables are given by: ***θ***= {*k*_prod_, *k*_bind_, *k*_unbind_, *kp, k*_dp_, *k*_deg_,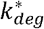 }. The cell state variables for the IGF/FoxO model are given by: ***θ***= { *k*_prod_, *k*_bind_, *k*_unbind_, *kp, k*_dp_, *k*_deg_,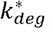, *k*_*ap*_, *k*_*adp*_, *k*_*in*_, *k*_*ef*_, *k*_*fb*_, *k*_*fdb*_,[*FoxO*]}. **D**. The estimated distribution of single cell mutual information values *p*_CeeMl_(*I*) for the EGF/EGFR pathway using maximum entropy estimation of *p*(***θ***)). The inset shows the input distribution *p*(*u*) corresponding to the maximum of the average *I*_Cee_ of *p*_CeeMl_(*I*) (blue), along with the input distribution corresponding to the channel capacity of *I*_*CSA*_ (green). **E**. Same as **D** for the IGF/FoxO pathway. We additionally show the experimentally estimated *p*_CeeMI_ (*I*)(pink).

### Experimental verification of *p*_CeeMI_ (*I*) using growth factor signaling networks

In cell populations, state variables that govern signaling dynamics such as protein levels (receptors, kinases, dephosphatases, etc.)^20, 21^ as well as effective rate constants such as endocytosis rates^39^, ligand binding rates^40^, etc. vary from cell to cell. Therefore, we expect the downstream phenotype of environmental sensing to be widely distributed as well. To experimentally verify the computational prediction of the distribution *p*_CeeMI_ (*I*) of sensing abilities, we need a system that allows us to approximate the cell state specific response distribution *p*(*x*|*u,**θ***). The IGF/FoxO pathway^35^ is an ideal system for these explorations for several reasons. First, following IGF stimulation, the transcription factor FoxO is pulled out of the nucleus. GFP-tagged variant of FoxO can be used to detect the dynamics of nuclear levels of FoxO at the single cell level^41^. Second, nuclear FoxO levels reach an approximate steady state within 30-45 minutes of constant IGF stimulation with FoxO levels decreasing with increasing IGF dose^41^. As a result, an approximate cell state specific distribution *p*(*x*|*u,**θ***) of steady state levels of FoxO can be obtained by stimulating the same cell with successive IGF levels. Finally, the biochemical interactions in the IGF/FoxO are well studied (Fig. 2C), allowing us to a build stochastic biochemical model based on previous computational efforts^42, 43^ that fits the single cell data accurately. Another system where *p*_CeeMI_ (*I*) can in principle be verified is the EGF/EGFR pathway (Fig. 2C). Here too, abundance of cell surface EGFR levels can be tracked in live cells following EGF stimulation^44^, allowing us to obtain cell-state specific response distribution *p*(*x*|*u,**θ***). Below, we show estimates of *p*_CeeMI_ (*I*)for both pathways and an experimental verification of our estimates for the IGF/FoxO pathway where live-cell imaging data was previously collected^41^.

The details of the calculations presented below can be found in SI Sections 3 and 4. Briefly, we first constructed stochastic biochemical models for the two pathways based on known interactions (Fig. 2C) and previous models^42, 43^. The output for the EGFR pathway was defined as the steady state levels of cell surface EGF receptors and the output for the IGF/FoxO pathway was defined as the steady state nuclear levels of the transcription factor FoxO. Using previously collected single cell data on the two pathways^37, 41, 43^ and our maximum entropy-based framework^37^, we estimated the distribution over parameters *p*(***θ***) for the two pathways (SI Section 3). Using these inferred distributions, and the model-predicted cell state specific response distribution *p*(*x*|*u,**θ***), we could compute *p*_CeeMI_ (*I*) for any specified input distribution *p*(*u*). We choose the support of the input distribution as the ligand concentrations used in the experimental setup. The estimates shown in Fig. 2D and Fig. 2E show *p*_CeeMI_ (*I*) corresponding to the input distribution *p*(*u*) that maximizes the conditional mutual information *I*_Cee_ (see Eq. 6). This input distribution is shown in the inset of Fig. 2D and 2E.

Similar to the illustrative example (Fig. 2A), there is a wide distribution of single cell sensing fidelities in real populations (Fig. 2D and Fig. 2E). Moreover, most cells are better sensors compared to the “average cell”, a cell whose response *p*(*x*|*u*) is averaged over cell state variability which was estimated to be ∼1 bit for both pathways. The cellular signaling fidelity skews towards the upper limit of 2 bits which corresponds to the logarithm of the number of inputs used in the experiment. Indeed, as seen in the insets of Fig. 2D and Fig. 2E, the input distribution corresponding to the maximum of the cell state agnostic mutual information *I*_CSA_ is concentrated on the lowest and the highest input for both pathways, indicating that cells may be able to detect only two input levels. In contrast, the input distribution corresponding to the maximum of *I*_Cee_ is close to uniform, suggesting that individual cells can in fact resolve different ligand levels.

To verify our computational estimate of *p*_CeeMI_ (*I*)for the IGF/FoxO pathway, we reanalyzed previously collected data wherein several cells were stimulated using successive IGF levels^41^. The details of our calculations can be found in SI Section 4. Briefly, the cells reached an approximate steady state within 60 minutes of each stimulation and nuclear FoxO levels measured between 60 to 90 minutes were used to approximate an experimental cell state specific response distribution *p*(*x*|*u,**θ***). The distribution *p*_CeeMI_ (*I*)was then obtained by maximizing the average mutual information *I*_Cee_ (averaged over all cells) with respect to the input distribution. As seen in Fig. 2D and Fig. 2E, the experimentally evaluated single cell signaling fidelities match closely with our computational estimates. Moreover, as predicted using our computational analysis, individual cells in the population were significantly better at sensing than what is implied by the maximum of *I*_CSA_. Indeed, the distribution of steady state FoxO levels were found to be well-resolved at the single cell level as well (Fig. 2E). Live cell imaging data for the EGFR pathway was not available and we leave it to future studies to validate our predictions.

Our calculations show that real cell populations comprise cells that have differing sensing fidelities, individual cells are significantly better at sensing their environment than what traditional estimates would indicate, and importantly, the CeeMI approach can accurately estimate the variation in signaling performances using readily collected experimental data and stochastic biochemical models. Notably, the variability in cell states and therefore the heterogeneity in sensing abilities is likely to be stable over time; the same cell’s FoxO responses to the same input were found to have significantly less variation compared to the variation within the population (SI Section 4).

### CeeMI identifies biochemical parameters that determine cells’ ability to sense their environment

Like other phenotypes^21^ and signaling dynamics^20, 30^, we expect that cells’ ability to sense their environment depends on their state variables. For example, cells with faster endocytosis rates^39^ may integrate environmental fluctuations with higher accuracy^43^. Or cells with higher receptor numbers^21^ will lead to a lowered relative noise in bound ligand concentration.

To systematically identify the variables that differentiate between cells’ ability to sense the environment, we quantify the joint distribution *p*_CeeMI_ (*I*, χ) of single cell signaling fidelity and biochemical state variables that take part in the signaling network. To test whether we can identify variables that are predictive of cellular fidelity, we estimated the joint distribution *p*_CeeMI_ (*I*, χ) for two variables that were experimentally accessible, response range of nuclear FoxO (Fig. 3 left, see inset) and total nuclear FoxO levels prior to IGF stimulation (Fig. 3 right, see inset). In both figures, the shaded regions show computational estimates of the joint probability distributions, and the red circles represent real cells. The green and the cyan trend lines represent computational and experimental binned averages.

**Figure 3.**
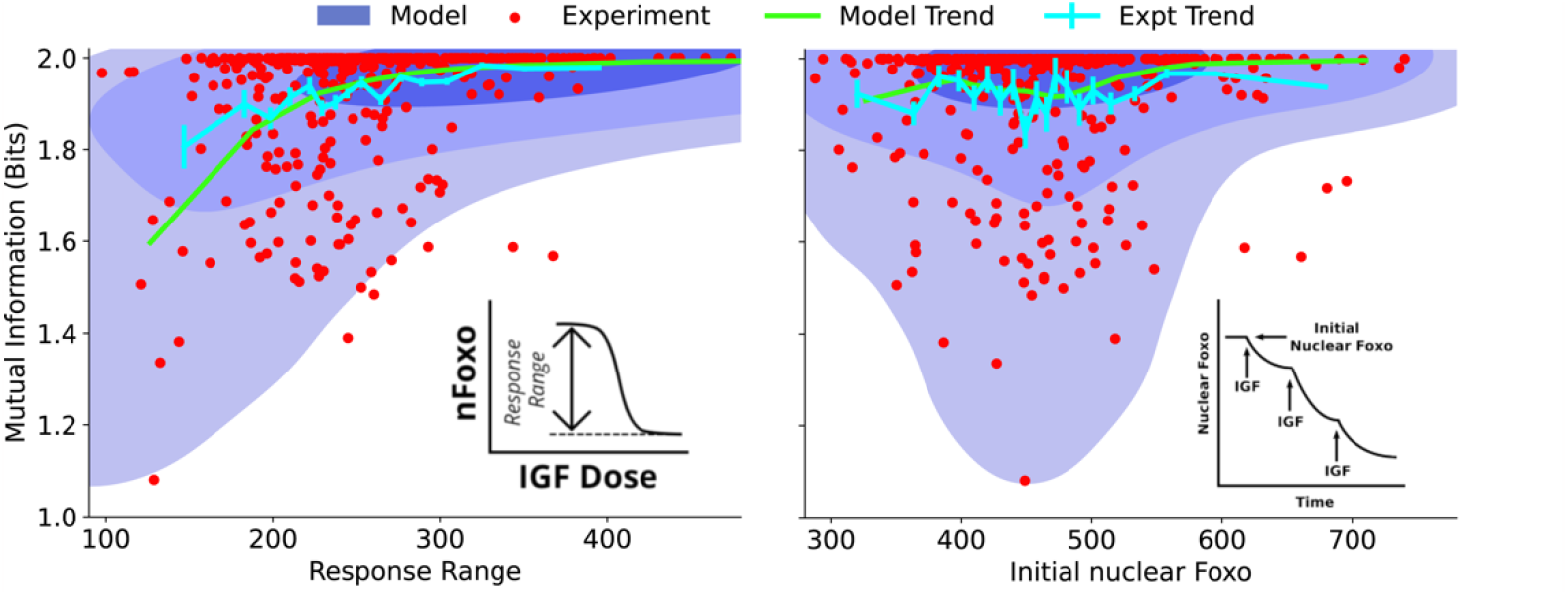
Dependence on cell state dependent mutual information on biochemical parameters. (left) The joint distribution *p*_CeeMI_ (*I*, χ) of cell state specific mutual information and biochemical parameter χ chosen to be the single cell response range of nuclear FoxO levels (x-axis, see inset for a cartoon). The shaded blue regions are model predictions, and the green line is the model average. The darker shades represent higher probabilities. The red dots represent experimental cells. The cyan line represents experimental averages. (right) same as (left) with biochemical parameter χ chosen to be steady state nuclear Foxo levels in the absence of stimulation. The contours represent 1% to 10%, 10% to 50%, and 50% to 100% of the total probability mass (from faint to dark shading).

One may expect that higher total nuclear FoxO levels could result in lower noise (coefficient of variation) and therefore better sensing abilities. However, we find that total nuclear FoxO levels only weakly correlate with cell state dependent mutual information (correlation coefficient *r* = 0.16 for computational estimates and *r* = 0.04 for experimental data). In contrast, cell state dependent mutual information depended strongly on the dynamic range of the response (correlation coefficient *r* = 0.53 for computational estimates and *r* = 0.29 for experimental data). Importantly, the model captured the observation that cells with a small response range had a variable sensing abilities while cells with a large response range all performed well in resolving extracellular IGF levels. Surprisingly, the total nuclear FoxO levels only weakly correlated with the cell specific mutual information. In SI Section 5, we show the predicted joint distributions *p*_CeeMI_ (*I*, χ) for several other biochemical variables that can potentially govern single cells’ response to extracellular IGF stimuli. This way, CeeMI can be used to systematically identify cell state variables that differentiate between good sensors and bad sensors of the environmental stimuli.

### Time scale of stochastic dynamics of cell states dictates the divergence between state-specific and state-agnostic sensing abilities

A key limitation of the presented analysis is the assumption that cell state variables remain approximately constant over the time scale of typical environmental fluctuations. While many cell state variables change slowly, remaining roughly constant over the life spans of cells and beyond^25, 26^, state changes may occur within the lifespan of a cell as well^45^. These dynamical changes can be accommodated easily in our calculations. Here, instead of fixing the cell state variables ***θ***,we can treat them as initial conditions and propagate them stochastically with their own dynamics. In the limit of very fast dynamics where individual cells rapidly transition cell states, we expect that the individuality of cells in a population vanishes and consequently cell state specific mutual information for each cell will agree with traditional cell state agnostic estimates of the channel capacity. In contrast, if the cell state dynamics are slow, our framework highlights differences between cells in the population.

To elucidate the role of cell state dynamics, we built a simple model of ligand-receptor system (SI Section 6, Fig. 4). Briefly, the model included production and degradation of the receptor mRNA, translation of the receptor protein from the mRNA, and ligand binding to the receptor (Fig.4A). The number of ligand-free receptors and the number mRNA molecules were together considered the state variables. We tuned the mRNA dynamics by keeping the average mRNA number constant while simultaneously changing mRNA production and degradation rates. The time scale of mRNA dynamics is denoted by τ.

**Figure 4.**
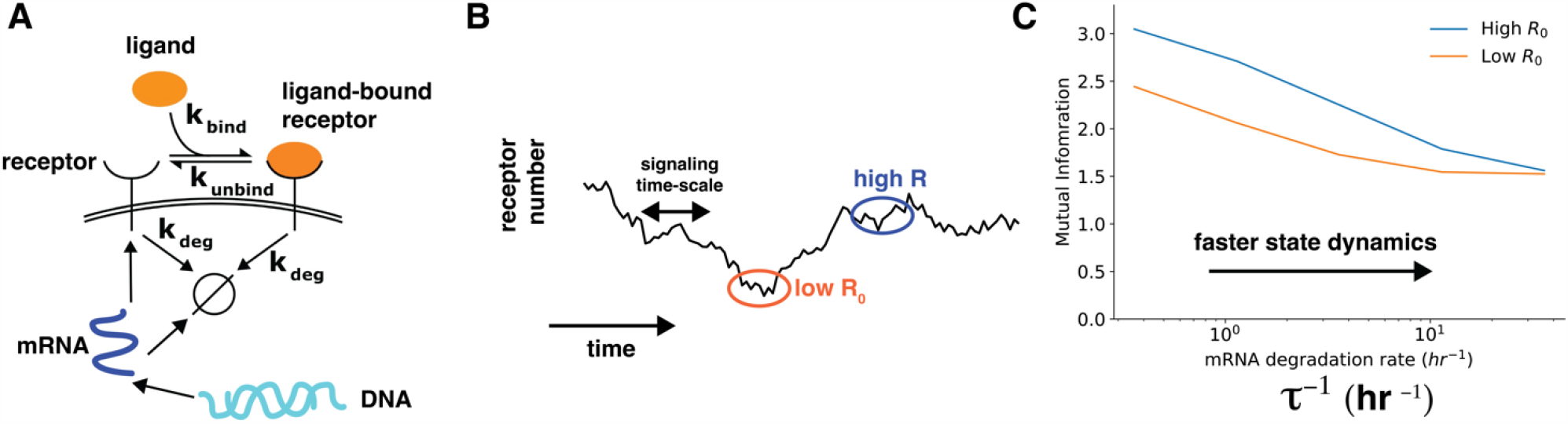
Cell state dynamics governs cell state conditioned mutual information. A. In a simple stochastic model, receptor mRNA is produced at a constant rate from the DNA and the translated into ligand-free receptors. The number of ligand-bound receptors after a short exposure to ligands is considered the output. B. A schematic showing dynamics of receptor numbers when mRNA dynamics are slower compared to signaling time scales. C. Conditioning on receptor numbers leads to differing abilities in sensing the environment when the time scale of mRNA dynamics is slow. In contrast, when the mRNA dynamics are fast (large τ^-1^), conditioning on cell state variables does not lead to difference in sensing abilities.

When the mRNA dynamics are slower than translation, ligand binding, and receptor degradation, individual cells are effectively frozen around a particular receptor number (Fig. 4B). The distribution of the number of ligand-bound receptors after a short-term ligand exposure reflects this frozen state. As a result, the cell-state conditioned mutual information depends strongly the cell state, with higher mutual information for cells with higher receptor numbers. In contrast, as the mRNA dynamics become faster, cells quickly lose the memory of cell state conditioning. As a result, the distribution of the number of ligand-bound receptors after a short-term ligand exposure reflects an ergodic averaging over the underlying mRNA dynamics. This averaging effectively washes out differences between cells (Fig. 4C).

This example shows that when cells states change at time scales slower than signaling dynamics, cells in a population can be identified by their cell state and their ability to sense the environment can differ from one another.

## Discussion

Cell populations are characterized by heterogeneity in cell state variables^18, 19^ that is responsible for important phenotypic differences and selective advantage for example in sensitivity to drugs^25, 26, 28^, response to chemotactic signals^46^, and following proliferation cues^29^. Therefore, it is reasonable to expect that cells’ ability to sense their environment depends on their state and is therefore variable across cells in a population. To quantify this heterogeneity, here, we developed a novel information theoretic perspective that allowed us to quantify the distribution *p*_CeeMI_ (*I*)of single cell sensing abilities using easily measurable single cell data and stochastic models of signaling networks. We also quantified the joint distribution *p*_CeeMI_ (*I*, χ) of cell specific sensing ability and biochemical cell state variable. Importantly, using two growth factor pathways, we showed that individual cells in real cell populations were much better at sensing their environment compared to what is implied by the traditional estimate of channel capacities of signaling networks. Typical single cell data are time stamped and do not give information about dynamics in the single cell. Moreover, these data do not provide information about dependence of cell-to-cell variability on cell state variables. Therefore, computational modeling of cellular trajectories from single cell data and stochastic models is essential in deciphering mechanistic origins of heterogeneity in cellular phenotypes.

The approach presented here will be useful in identifying bottlenecks in signal transduction. Many cellular phenotypes such as chemotaxis^7, 8^ and cell proliferation^47^ exhibit a weak correlation between cellular outputs (e.g., directional alignment with chemical gradients) with the input (e.g., gradient strength), resulting in a low channel capacity even for individual cells. In such cases, it is important to understand where exactly along the information transduction pathway is the information about the gradient is lost. If traditional calculations are pursued, for example, for movement of mammalian cells under growth factor gradients^7, 31^, one may conclude that the information loss likely happens right at the receptor level (Fig. 2D). In contrast, CeeMI will allow us to disentangle the effect of cell state heterogeneity and noisy cellular response to precisely pinpoint intracellular signaling nodes that are responsible for signal corruption.

How do we contrast our results with previous low estimates of channel capacities? There is extensive population heterogeneity in cell state variables and this heterogeneity often is stable over time. Nonetheless, this heterogeneity arises from stochasticity in underlying processes including dynamics of epigenetic transitions and gene expression. When one specifies state variables^19^ ***θ***, one effectively conditions on the stochastic trajectory of the underlying dynamics to a specific narrow window (Fig. 4). This extra conditioning narrows the cells’ response distributions, revealing the heterogeneous nature of cell populations^19^.

In summary, we showed that like other phenotypes, the ability to sense the environment is itself heterogeneously distributed in a cell population. Moreover, we also showed that when conditioned on cell state variables, mammalian cells appear to be significantly better at sensing their environment than what traditional mutual information calculation suggests. Finally, we showed that we could identify cell state variables that made some cells better sensors compared to others. We believe that CeeMI will be an important framework in quantifying fidelity of input/output relationships in heterogeneous cell populations.

## Acknowledgments

AG, HA, and PD are supported by NIGMS grant R35GM142547. The authors would like to thank Andre Levchenko for useful discussions.

## Notes

### Competing Interest Statement

The authors have declared no competing interest.

### Summary of Updates

We have revised the manuscript in response to comments from reviewers at eLife.

