## Supplementary file for "The ability to sense the environment is heterogeneously distributed in cell populations"

Scripts used to generate mutual information results as well the experimental data used to train our models can be obtained at: <https://github.com/adgoetz186/Cell_signalling_information>. Script for maximum entropy inference of cellular parameters can be obtained at: <https://github.com/hodaakl/MaxEnt>.

The supplementary materials are organized as follows: Section 1 describes basic information theoretical concepts and our numerical approach to estimate mutual information and channel capacity. Section 2 describes the *in-silico toy* model. Section 3 describes the maximum entropy approach. Section 4 describes numerical methods for quantification of mutual information and channel capacity using live cell imagining data. Section 5 shows the joint distribution of mutual information and biochemical parameters. Section 6 describes the methods to examine the role of cell state dynamics on sensing ability.

1. **Information theory primer**

Here, we give a brief description of information theoretic quantities used in the manuscript. The readers are referred to a textbook^1^ for a detailed discussion.

A communication channel (for example, a signaling network) is an input/output relationship between two random variables: the input $U$ (say, a ligand concentration) and the response $R$ (e.g. levels of some intracellular protein). The mutual information (MI) between $U$ and $R$ is the reduction in the uncertainty about $U$ due to the access of the outcome of $R$. The MI is defined as

$$\begin{aligned} I\equiv I\left( U;R \right)=\sum_{r\in R,u\in U} p\left( r|u \right)p\left( u \right)\log_{2} \frac{p\left( r | u \right)}{\sum_{u^{'}} p\left( r | u^{'} \right)p\left( u^{'} \right)}\#\left( S1 \right) \end{aligned}$$

where the choice of two in the base of the logarithm gives the value of information in bits. The summation is replaced by integrals when considering continuous random variables. In the context of cell signaling, we will use the notation $I_{CSA}$ (instead of $I$ in Eq. S1) to denote the cell state agnostic mutual information, which is the mutual information between the response distribution $p\left( r | u \right)$ and the input distribution $p\left( u \right)$.

We also consider a more nuanced situation where instead of a single channel, we have a family of channels whose states are characterized by a random variable $\Theta$. The channel state $\boldsymbol{\theta}\in\Theta$ uniquely determines the probabilistic response relationship $p\left( r|u,\boldsymbol{\theta} \right)$ between $U$ and $R$ when $\Theta$ is fixed. Similar to Eq. S1, we can define the MI between $U$ and $R$ conditioned for a specific realization $\boldsymbol{\theta}\in\Theta$

$$\begin{aligned} I\left( \boldsymbol{\theta} \right)=\sum_{r\in R,u\in U} p\left( r|u,\boldsymbol{\theta} \right)p\left( u \right)\log_{2} \frac{p\left( r | u,\boldsymbol{\theta} \right)}{\sum_{u^{'}} p\left( r | u^{'},\boldsymbol{\theta} \right)p\left( u^{'} \right)}\#\left( S2 \right) \end{aligned}$$

The average of $I(\boldsymbol{\theta})$ over $p(\boldsymbol{\theta})$ is traditionally known as the conditional mutual information^1^. We define the **Ce**ll stat**e** dependent mutual information $I_{Cee}:$

$$\begin{aligned} I_{Cee}\equiv I\left( U;R|\Theta\right)=\sum_{\boldsymbol{\theta}\in\Theta} I\left( \boldsymbol{\theta} \right)p\left( \boldsymbol{\theta} \right).\#\left( S3 \right) \end{aligned}$$

The channel capacity (CC) is a measure of the optimal performance of the signaling network with respect to the distribution of $U$. It can be defined for both $I$ and $I_{Cee}$ as follows,

$$\begin{aligned} CC_{I}=\max_{p\left( u \right)} I\left( U;R \right) \mathrm{or}CC_{I_{Cee}}=\max_{p\left( u \right)} I\left( U;R | \Theta\right)\#\left( S4 \right) \end{aligned}$$

The difference between $I$ and $I_{Cee}$ is called the interaction information $I(U;R;\Theta)$. The interaction information can be simplified as

$$\begin{aligned} I\left( U;R;\Theta\right)=I\left( U;R \right)-I\left( U;R|\Theta\right)=I\left( U;\Theta\right)-I\left( U;\Theta|R \right)=-I\left( U;\Theta|R \right)\leq0\#\left( S5 \right) \end{aligned}$$

where $I\left( U;\Theta\right)=0$ follows from the statistical independence of the input signal $U$ and channel state $\Theta$. Eq. S5 shows that $I_{Cee}>I$ as long as $I\left( U;\Theta\right)=0$. We note that this may not be true in general. However, it is true in the context of cellular signaling networks where the inputs are chosen by the experimentalists while the cell states are an inherent property of the cell population. It is trivial to extend this argument to the channel capacities of both terms.

**1.1 Numerical estimation of mutual information**

Evaluating the mutual information between an input and an output (Eq. S1 and Eq. S2) requires numerical integration over the input and the output distribution. We limit the input distributions to a finite support, specifically to the ligand concentrations that were used in the experimental set up. These are $L=[0,0.0078,0.01,0.03,0.06,0.125,0.25,0.5,1,100]$ ng/mL for the EGF/EGFR pathway and $L=[0, 17.5, 37.5, 125]$ pM for the IGF/FoxO pathway.

The numerical integration in Eq. S1 and Eq. S2 requires summing over probabilities of all possible responses and inputs. When the response distributions are approximated using histograms (either from experimental data or from Markov chain Monte Carlo simulations of a model), the summation can be error prone. To avoid this, we assume that the responses are distributed according to a gamma distribution (see Fig. S4 below). The gamma distribution was chosen here and in several other places because it has been shown to accurately approximate real distributions of protein/mRNA abundances^2^.

To speed up the calculations, inspired by previous approaches^3^, we use a binning strategy. Specifically, we bin the response distribution using a constant bin width. The bin width was chosen to be equal to 5% of the smallest inter-quartile range across response distributions corresponding to all considered inputs. The binning procedure requires truncating the response distributions to a finite support. We ensured that our binning captured at least 99.95% of the entire mass of the distribution. The same strategy was used to obtain $I_{CSA}$ in Eq. S1 and $I(\boldsymbol{\theta})$ in Eq. S2. The only difference in computing $I(\boldsymbol{\theta})$ compared to $I_{CSA}$ was that the response distribution $p(r|u,\boldsymbol{\theta})$ was obtained computationally using a stochastic differential equation model with network parameters fixed at $\boldsymbol{\theta}$. Samples from the joint distribution $p_{CeeMI}(I,\chi)$ where $\chi$ is a cell state variable were obtained by sampling cell state variables $\boldsymbol{\theta}$ from $p(\boldsymbol{\theta)}$ (see section 3) and simultaneously evaluating $I(\boldsymbol{\theta})$ and $\chi(\boldsymbol{\theta})$.

Once the mutual information could be obtained numerically for a given input distribution $p(u)$, the corresponding channel capacity, the maximum of the mutual information over all possible input distributions, can be obtained by solving the following optimization problem:

$$\begin{aligned} \max_{p\left( u \right)} I_{CSA} s.t. \sum p\left( u \right)=1 \mathrm{and}p\left( u \right)\geq0\#\left( S6 \right) \end{aligned}$$

The optimization problem was solved via a trust region constrained algorithm^4^ using the SciPy optimization library^5^. When the distribution over cell state variables $p\left( \boldsymbol{\theta} \right)$ is available, we can estimate $I_{Cee}$ as the average $\left\langle I\left( \boldsymbol{\theta} \right) \right\rangle_{\boldsymbol{\theta}}$ (Eq. S3). The optimum value of $I_{Cee}$ over all input distributions serves as a cell state dependent analogue to the channel capacity of $I_{CSA}$ and is obtained by solve a similar optimization problem:

$$\begin{aligned} \max_{p\left( u \right)} I_{Cee} s.t. \sum p\left( u \right)=1 \mathrm{and}p\left( u \right)\geq0\#(S7) \end{aligned}$$

The procedure needed to evaluate single cell mutual information values using live cell imaging data on the IGF/FoxO pathway is described in section 4.

2. **Description of the *in-silico* toy model**

The *in-silico* cell receptor/ligand system comprises two components (unbound receptors and bound receptors) and five reactions (Fig. S1).

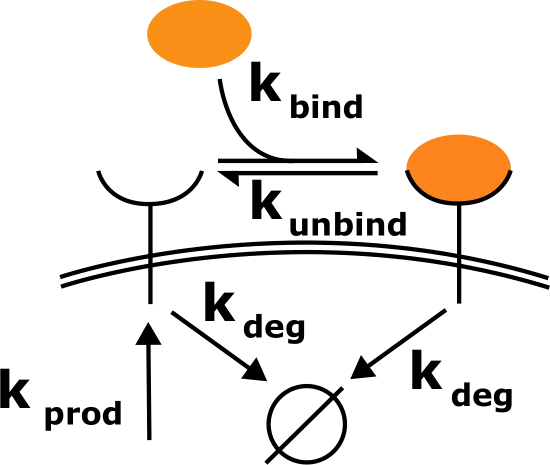

**Figure S1. Schematic of the toy model.** Cell surface receptors bind to ligand (orange oval). Ligand free receptors and ligand bound receptors are shuttled between the plasma membrane and the cytoplasm.

In the model, receptors are constantly shuttled to the membrane and removed from the membrane and degraded. The extracellular ligand (concentration denoted by $L$) binds to cell surface receptors. We assume that the ligand concentration is kept constant in the environment. Steady state abundance of ligand bound receptor (denoted by $B$) is taken as the output of the system. At steady state, the mean field equations for the average species level are:

$$\begin{aligned} k_{\mathrm{prod}}-k_{\mathrm{bind}}LR+k_{\mathrm{unbind}}B-k_{\deg}R=0\#\left( S8 \right) \end{aligned}$$

$$\begin{aligned} k_{\mathrm{bind}}LR-k_{\mathrm{unbind}}B-k_{\deg}B=0\#\left( S9 \right) \end{aligned}$$

Solving for steady state, at the single cell level, the mean number of bound receptors is given by

$$\begin{aligned} \mu_{B}=R_{0}\frac{Lk_{\mathrm{bind}}}{Lk_{\mathrm{bind}}+k_{\deg}+k_{\mathrm{unbind}}}.\#\left( S10 \right) \end{aligned}$$

In Eq. S10, $k_{\mathrm{bind}}$ is the ligand binding rate, $k_{\mathrm{unbind}}$ is the ligand unbinding rate, $k_{\deg}$ is the degradation rate, and $R_{0}$ is the average value of the receptor level in the absence of the ligand. The bound receptor levels are Poisson distributed with a mean given by Eq. S10 (Fig. S2A)

Using this toy network, we created two *in silico* cell populations. In both populations, we fixed $k_{\mathrm{bind}}=1 {sec}^{-1}a.u.^{-1}$ and $k_{\mathrm{unbind}}=10 {sec}^{-1}$. In the first population, every parameter was kept constant across cells except for the cell surface receptor levels $R_{0}$. In the second population, every parameter was kept constant across cells except for the receptor degradation rate $k_{\deg}$. In the first population (when $R_{0}$ was varied) we fixed $k_{\deg}=5{sec}^{-1}$. In the second population (when $k_{\deg}$ was varied), we fixed $R_{0}=50 molecules/cell$. The parameter that varied across cells was assumed to be distributed according to a gamma distribution. For the two populations, we kept fixed mean value of the variable parameter to be ${\langle R}_{0}\rangle=500 molecules/cell$ and ${\langle k}_{\deg}\rangle=5 {sec}^{-1}$. We varied coefficients of variation for both populations between $CV={10}^{-1.5}$ and $CV={10}^{-0.5}.$

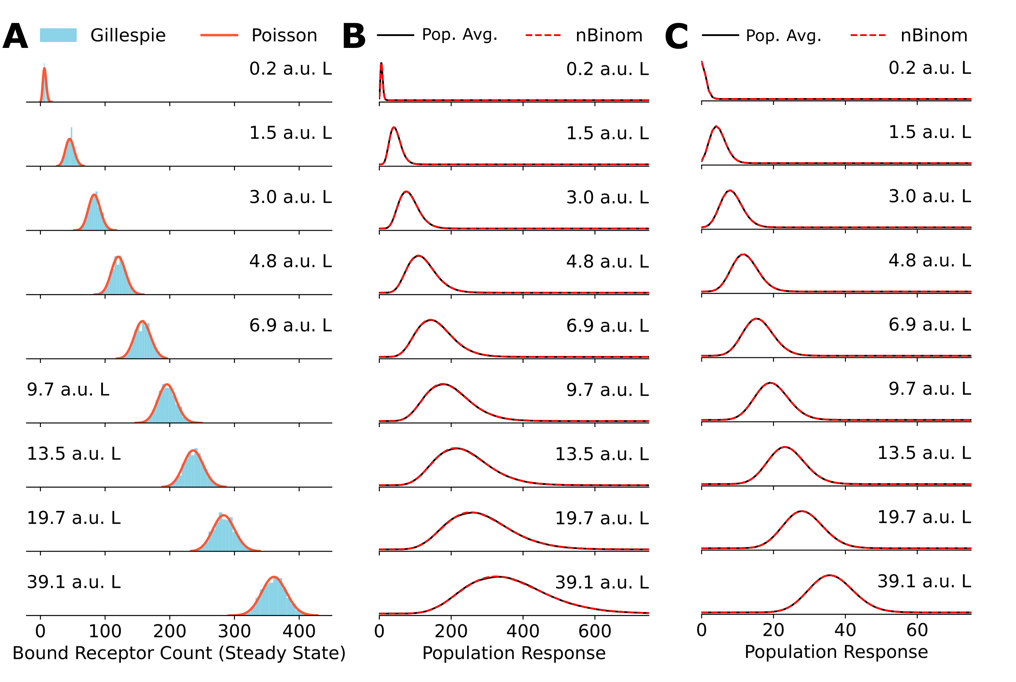

**Figure S2. A**. Comparison between explicit simulations of the toy network using Gillespie’s algorithm and the Poisson distribution. **B**. A comparison between the negative binomial distribution assumption and population average distribution of receptor levels obtained by averaging over a gamma distribution for the cell state variable $R_{0}.$ **C**. Same as in B for cell state variable $k_{\deg}.$

**2.1 Obtaining** $\boldsymbol{I}_{\boldsymbol{CSA}}$ **for the toy network**

Eq. 1 of main text shows that calculation of MI in a cell state agnostic manner requires the knowledge of the cell state averaged response distribution $p(B|L)$ and the distribution of inputs $p(L)$. In our calculations, we restrict $p(L)$ to be a discrete version of the gamma distribution obtained by equal percentile binning of a gamma distribution. The mean of the gamma distribution was taken to be 10, the coefficient of variation 1, and the number of bins was 25. The discretization step was not essential but was taken to simplify the calculations by making all variables discrete.

We assumed that $p(B|L)$ which is obtained by averaging over the cell state variables was distributed as a negative binomial distribution (Fig. S2B and C). We estimated its first two moments by numerically averaging the first two moments of the single cell response distribution $p(B|L,\boldsymbol{\theta})$ (Poisson distribution with mean given by Eq. S10) according to the population distribution $p(\boldsymbol{\theta})$ of the variable parameter(s) $\boldsymbol{\theta}$ ($\boldsymbol{\theta}\equiv R_{0}$ for the first cell population and $\boldsymbol{\theta}\equiv k_{\deg}$ for the second cell population). As mentioned above, $p(\boldsymbol{\theta})$ was assumed to be a gamma distribution whose coefficient was systematically varied. Using the first two moments, we inferred the parameters for the negative binomial distribution.

Once the population level response $p(B|L)$ is obtained and the input distribution $p(L)$ is fixed, we can calculate $I_{CSA}$ using Eq. 1. The values of $I_{CSA}$ for the population with variable $R_{0}$ are given in Fig. 2C of the main text while those for the population with variable $k_{\deg}$ are given in Fig. S3.

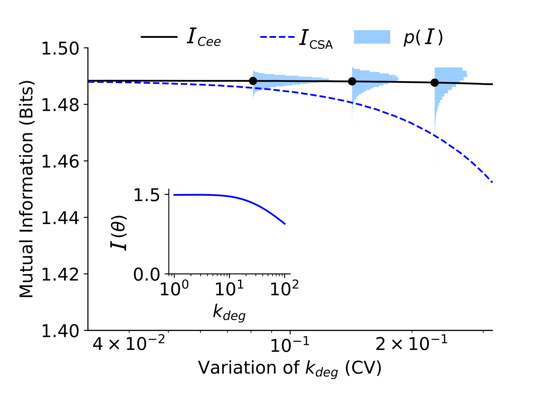

**Figure S3.** Values of $I_{CSA}$ (Eq. S1), $I_{Cee}$ (Eq. S3), and $p_{CeeMI}(I)$ (y-axis) as a function of the degree of variability (coefficient of variation) in $k_{\deg}$. The inset shows $I(\boldsymbol{\theta})$ (Eq. S2) as a function of $k_{\deg}$.

**2.2 Obtaining** $\boldsymbol{p}_{\boldsymbol{CeeMI}}\boldsymbol{(I)}$ **and** $\boldsymbol{I}_{\boldsymbol{Cee}}$ **for the toy receptor network**

As indicated by Eq. 4 and Eq. 6 of the main text, calculation of the distribution of cell state dependent mutual information values requires the cell state specific response distribution $p(B|L,\boldsymbol{\theta})$ and the distribution $p(\boldsymbol{\theta})$ of cell states. In the toy model, $p(B|L,\boldsymbol{\theta})$ is modeled as a Poisson distribution with a mean given by Eq. S8 and $p(\boldsymbol{\theta})$ is assumed to be gamma distributed in the variable parameter.

Using $p(B|L,\boldsymbol{\theta})$ and gamma distributed input distribution $p(L)$ we obtain cell state specific mutual information $I\boldsymbol{(\theta)}$ using Eq. S2. We find $p_{CeeMI}(I)$ by sampling multiple values of $\boldsymbol{\theta}$ ($R_{0}$ for the first population and $k_{\deg}$ for the second population). $I_{Cee}$ is simply the average of $p_{CeeMI}(I)$. $I_{Cee}$ and $p_{CeeMI}(I)$ for the population with variable $R_{0}$ are given in Fig. 2C of the main text. Fig. S3 show the same for the population with variable $k_{\deg}$.

3. **Maximum entropy inference of cell state variability**

Using experimentally collected single cell data on heterogeneity in protein abundances, we estimate the distribution over cell state variables (biochemical parameters) using the maximum entropy approach. Below, we first describe the data that was used in our analysis. Next, we briefly discuss the maximum entropy approach.

3.1 **EGF/EGFR pathway, data and model**

We used previously collected single cell data on cell surface EGFR levels^6, 7^. Briefly, MCF10A cells were stimulated with 10 different extracellular EGF levels ranging between 0ng/mL to 100 ng/mL. Cell surface EGFR levels were measured in $\sim7000$ cells for each ligand concentration after 3 hours of continuous EGF stimulation. The data was measured in arbitrary fluorescence units. To convert the data to the units of number of receptors per cell, we used a mean number of cell surface receptors $R=2.5\times{10}^{5}$ for MCF10A cells^8^. The population mean of the experimentally measured steady state receptor count distribution in the absence of the ligand was matched to this number. Receptor count data at every other ligand concentration was scaled appropriately.

Using a previously validated model^6, 7^, we constructed a simplified model of the EGF/EGFR pathway. Specifically, we incorporated ligand binding to receptor, receptor activation, and preferential endocytosis of activated receptors. To keep our model simple, we did not incorporate receptor dimerization and oligomerization following ligand binding. Notably, evidence suggests that oligomers may be pre-formed in the absence of the ligand as well, which could make them effective monomers. Finally, we assumed that the extracellular ligand concentration was kept constant. The model was represented by the following reaction network.

$$\begin{aligned} \phi\underset{\to}{k_{\mathrm{prod}}}R\#\left( S11 \right) \end{aligned}$$

$$\begin{aligned} R\underset{\to}{Lk_{\mathrm{bind}}}B, B\underset{\to}{k_{\mathrm{unbind}}}L+R\#\left( S12 \right) \end{aligned}$$

$$\begin{aligned} B\underset{\to}{k_{p}}P, P\underset{\to}{k_{\mathrm{dp}}}B\#\left( S13 \right) \end{aligned}$$

$$\begin{aligned} R\underset{\to}{k_{\deg}}\phi, B\underset{\to}{k_{\deg}}\phi, P\underset{\to}{k_{\deg}^{*}}\phi\#\left( S14 \right) \end{aligned}$$

In Eqs. S11-S14, $R$ is the level of free receptor, $B$ is the level of ligand bound receptor, and $P$ is the level of phosphorylated receptor. The total number of receptors is given by $R_{T}=R+B+P.$ Cell state variables $\boldsymbol{\theta}$ comprised $\boldsymbol{\theta\equiv}\left\{ k_{prod}, k_{bind}, k_{unbind}, k_{p}, k_{dp}, k_{deg}, k_{deg}^{*} \right\}\boldsymbol{.}$

**3.2 IGF/FoxO pathway, data and model**

We used previously collected single cell data on nuclear FoxO levels following continuous IGF stimulation in HeLa cells^9^. Briefly, cells were continuously stimulated with IGF, nuclear FoxO levels were measured using GFP-tagged FoxO using live cell imaging every 3 minutes for 90 minutes. To convert the arbitrary fluorescence units to units of copies of FoxO per cell, we first removed the background fluorescence intensity. Then, we used the previously estimated total FoxO levels in HeLa cells^10^ ($\sim710$ molecules/cell) and the nuclear to cytoplasmic ratio in the absence of stimulation^11^ ($2/3$ of the total in the nucleus). There was a small disagreement in mean nuclear FoxO levels in the absence of stimulation across different experiments. To remove this artifact and to start all experiments with the average nuclear FoxO levels, we offset individual experiments such that the mean nuclear FoxO was identical across all experiments.

Similar to the EGF/EGFR pathway, using a previously validated model^11^, we constructed a simplified model of the IGF/FoxO pathway. Specifically, we incorporated ligand binding to IGF receptor, receptor activation, activation of Akt, and Akt-driven phosphorylation of FoxO. Phosphorylated FoxO was prohibited from entering the nucleus. Finally, we assumed that the extracellular ligand concentration was kept constant. The model was represented by the following reaction network.

$$\begin{aligned} \phi\underset{\to}{k_{\mathrm{prod}}}R\#\left( S15 \right) \end{aligned}$$

$$\begin{aligned} R\underset{\to}{Lk_{\mathrm{bind}}}B, B\underset{\to}{k_{\mathrm{unbind}}}L+R\#\left( S16 \right) \end{aligned}$$

$$\begin{aligned} B\underset{\to}{k_{p}}P, P\underset{\to}{k_{\mathrm{dp}}}B\#\left( S17 \right) \end{aligned}$$

$$\begin{aligned} R\underset{\to}{k_{\deg}}\phi, B\underset{\to}{k_{\deg}}\phi, P\underset{\to}{k_{\deg}}\phi\#\left( S18 \right) \end{aligned}$$

$$\begin{aligned} Akt\underset{\to}{{P \cdot k}_{\mathrm{ap}}}pAkt, pAkt\underset{\to}{k_{\mathrm{adp}}}Akt\#\left( S19 \right) \end{aligned}$$

$$\begin{aligned} FoxO_{c}\underset{\to}{k_{in}}FoxO_{n}, FoxO_{n}\underset{\to}{k_{\mathrm{ef}}}FoxO_{c}\#\left( S20 \right) \end{aligned}$$

$$\begin{aligned} FoxO_{c}\underset{\to}{{pAkt\cdot k}_{fp}}pFoxO_{c}, pFoxO_{c}\underset{\to}{k_{\mathrm{fdp}}}FoxO_{c}\#\left( S21 \right) \end{aligned}$$

In Eqs. S15-S21, $R$ is the level of free IGF receptor, $B$ is the level of ligand bound receptor, and $P$ is the level of phosphorylated receptor. $pAkt$ is phosphorylated Akt, $pFoxO_{c}$ is cytoplasmic phosphorylated FoxO, $FoxO_{c}$ is cytoplasmic unphosphorylated FoxO, and $FoxO_{n}$ is nuclear unphosphorylated FoxO. Cell state variables $\boldsymbol{\theta}$ comprised $\boldsymbol{\theta\equiv}\left\{ k_{prod}, k_{bind}, k_{unbind}, k_{p}, k_{dp}, k_{deg}, k_{deg}^{*}, \left[ Akt \right], k_{ap}, k_{adp}, k_{in}, k_{ef}, k_{fp}, k_{fdp}, [FoxO] \right\}\boldsymbol{.}$

**3.3 Inference of model parameters**

We assume that cells in a population can be assigned a cell specific state denoted by a state vector $\boldsymbol{\theta}$ that comprises biochemical parameters relevant to the modeled signaling network. The population variability in cell state parameters is represented by the joint probability density $p\left( \boldsymbol{\theta} \right)\boldsymbol{.}$ Typically, $p\left( \boldsymbol{\theta} \right)$ is not experimentally accessible. Therefore, we infer it using a previously developed technique called MEDIRIAN (maximum entropy-based framework for inference of heterogeneity in dynamics of signaling networks)^6^. MERIDIAN infers the maximum entropy distribution $p\left( \boldsymbol{\theta} \right)$ that reproduces a set of averages computed from experimental single cell measurements.

MERIDIAN requires a mechanistic model of the signaling network that can predict cell's response to extracellular perturbation (e.g. ligand) and user-specified population averages computed from experimental data. For the EGF/EGFR and IGF/FoxO networks, we used stochastic biochemical models described by Eq. S11-S14, and S15-S21 respectively. We use the moment closure approximation^12^ to approximate the single cell distributions using the first two moments (see below). The differential equations for the pathways can be found on the github.

Now, we briefly describe the MERIDIAN approach**.** The entropy of any distribution $p\boldsymbol{(\theta)}$ is given by

$$\begin{aligned} S=-\int\boldsymbol{p}\left( \boldsymbol{\theta} \right)\log\boldsymbol{p}\left( \boldsymbol{\theta} \right)d\boldsymbol{\theta.}\#\left( S22 \right) \end{aligned}$$

In MERIDIAN, we find $p\boldsymbol{(\theta)}$ that maximizes $S$ while requiring it to reproduce a set of average constraints evaluated using experimental data. Following Dixit et al.^6^, entropy maximized $p\boldsymbol{(\theta)}$ is given by the Gibbs Boltzmann distribution

$$\begin{aligned} p\left( \boldsymbol{\theta} \right)\boldsymbol{=}\frac{1}{\Omega}\exp\left( -\sum_{m} \lambda_{m}\psi_{m}\left( \boldsymbol{\theta} \right) \right)\#\left( S23 \right) \end{aligned}$$

where $\lambda_{m}$ are the Lagrange multiplier corresponding to the $m^{\mathrm{th}}$ constraint, $\psi_{m}\left( \boldsymbol{\theta} \right)$ is the quantity whose average is constrained, and a $\Omega$ is the normalization constant.

Given a set of constraints (see below), the Lagrange multipliers can be numerically tuned such that the predictions from the distribution $p\left( \boldsymbol{\theta} \right)$ match their experimental value. We optimize the Lagrange multipliers using gradient-based search (Eq. S24) using the ADAM algorithm^13^. The gradients for minimizing a Lagrangian cost function are given by^6^

$$\begin{aligned} L=\log\Omega+\sum_{m} \lambda_{m}R_{m} \\ \frac{\partial L}{\partial\lambda_{m}}=R_{m}-\left\langle\psi_{m}\left( \boldsymbol{\theta} \right) \right\rangle_{\boldsymbol{\theta}}.\#\left( S24 \right) \end{aligned}$$

In Eq. S24, $\left\langle\psi_{m}\left( \boldsymbol{\theta} \right) \right\rangle_{\boldsymbol{\theta}}$ denotes the ensemble average computed using $p\left( \boldsymbol{\theta} \right)$ and $R_{m}$ are the corresponding measurements. We stop the iterative procedure when the mean absolute relative error $\frac{1}{M}\sum_{m} \frac{\left| R_{m}-\left\langle\psi_{m}\left( \boldsymbol{\theta} \right) \right\rangle_{\boldsymbol{\theta}} \right|}{R_{m}}$ reaches a predefined value.

The predictions from the Max Ent model depends on the choice of the experimental constraints. To ensure that the constraints represent the entire range of single cell behaviors, we opt for percentile constraints. Specifically, for experimentally collected single cell data for a given condition (ligand dose, time point, etc.), we first approximate the single cell histogram using a gamma distribution. Then, we identify abundances that represent 10^th^ to 90^th^ percentiles of this distribution. The fraction of cells belonging to each of these percentile windows is exactly 10%. These become our experimental constraints $R_{em}=0.1$. Here, $"e"$ denotes the experimental condition (ligand dose, time point, etc.) and $m\in[1,10]$ denotes the percentile window.

The corresponding model predictions of the fraction of cells in a given percentile window $\left\langle\psi_{em}\left( \boldsymbol{\theta} \right) \right\rangle_{\boldsymbol{\theta}}$ are given by

$$\begin{aligned} \left\langle\psi_{em}\left( \boldsymbol{\theta} \right) \right\rangle_{\boldsymbol{\theta}}=\int p\left( \boldsymbol{\theta} \right)\psi_{em}\left( \boldsymbol{\theta} \right)d\boldsymbol{\theta}\#\left( S25 \right) \end{aligned}$$

where

$$\begin{aligned} \psi_{em}\left( \boldsymbol{\theta} \right)=\int_{l_{em}}^{u_{em}} p_{e}\left( r|\boldsymbol{\theta} \right)dr.\#\left( S26 \right) \end{aligned}$$

In Eq. S26, $p_{e}\left( r|\boldsymbol{\theta} \right)$ is the model predicted single cell distribution of responses (surface EGFR levels or nuclear FoxO levels) for experimental condition $e$. The integration bounds $l_{em}$ and $u_{em}$ represent the lower and upper bounds of the $m^{\mathrm{th}}$ percentile window. The distribution $p_{e}\left( r|\boldsymbol{\theta} \right)$ in principle can be approximated by several runs of an explicit simulation using Gillespie’s algorithm^14^. This may prove to be computationally expensive, especially when sampling through multiple parameter sets $\boldsymbol{\theta}$. Therefore, we resort to moment closure techniques^12^ to approximate $p_{e}\left( r|\boldsymbol{\theta} \right)$ as a gamma distribution. We used a previously developed package called MOCA (moment closure analysis)^14^. We used a Gaussian moment closure to obtain the first and the second moments of the distributions. The moment closure approximation was quite accurate compared to the explicit Gillespie simulation and allowed us to rapidly predict single cell response distributions without performing multiple calculations (Fig. S4)

The averages in Eq. S25 cannot be computed analytically. We therefore resort to Markov chain Monte Carlo techniques to approximate them. Briefly, for a fixed set of Lagrange multipliers, we start 150 parallel MCMC chains in the parameter space with a starting point chosen randomly from the previous iteration. Each step in the MCMC calculation attempted to change between 1 to 5 parameters. The step size for the change was a uniform random number whose maximum was 10% of the parameter bounds for individual parameters. Each MCMC chain was run for approximately for $\sim{10}^{4}$ steps for the EGF/EGFR pathway and $\sim2.5\times{10}^{5}$ for the IGF/FoxO pathway. The first 1000 steps were discarded, and parameter sets were stored every 50^th^ step after that.

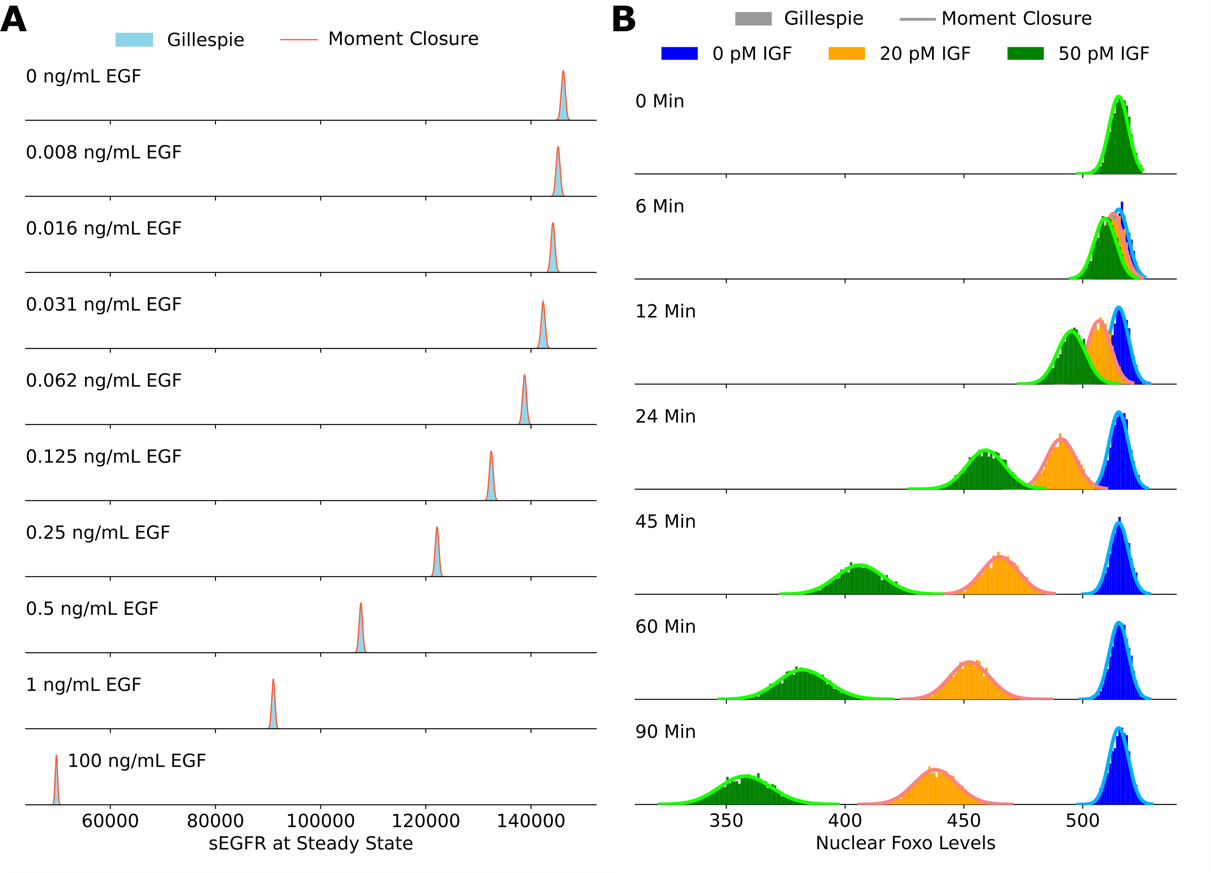

**Figure S4.** Validating the accuracy of the moment closure approximation against histograms obtained using the Gillespie algorithm. The steady state cell surface EGFR levels are shown on the left and the steady state nuclear FoxO levels are shown on the right.

We used ADAM to optimize the Lagrange multipliers (Fig. S5). The hyperparameters for ADAM were as follows: the exponential decay rate for the first moment estimates was set to 0.8. The exponential decay rate for the second-moment estimates was set to 0.999, the step size was 0.1. This procedure led to a decrease in the relative error. Using the final set of Lagrange multipliers, we sampled parameter sets that represented an *in-silico* cell population. This population was used for further analysis (see below).

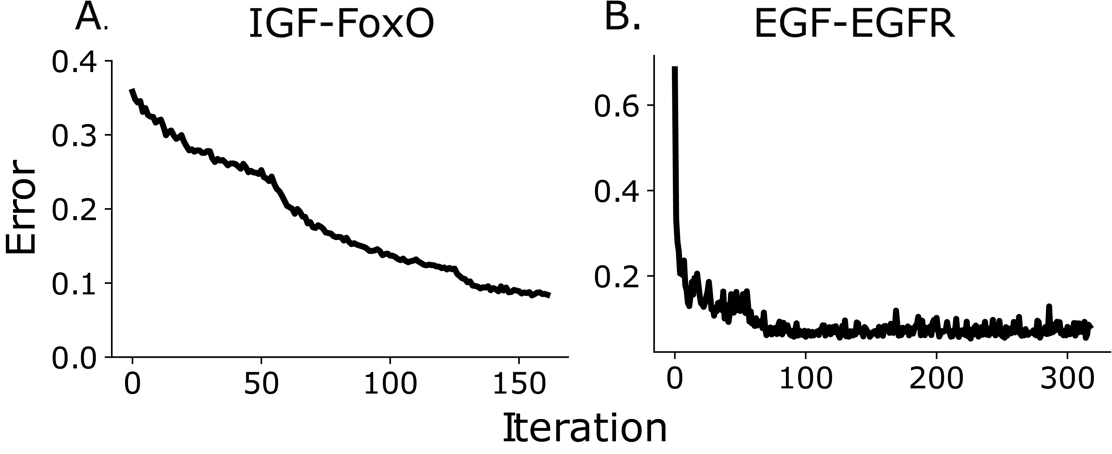

**Figure S5.** Mean absolute relative error between predicted bin fractions and estimated bin fractions as a function of iteration number for the IGF/FoxO and the EGF/EGFR pathway.

4. **Numerical quantification of mutual information and channel capacity using live cell imaging data**

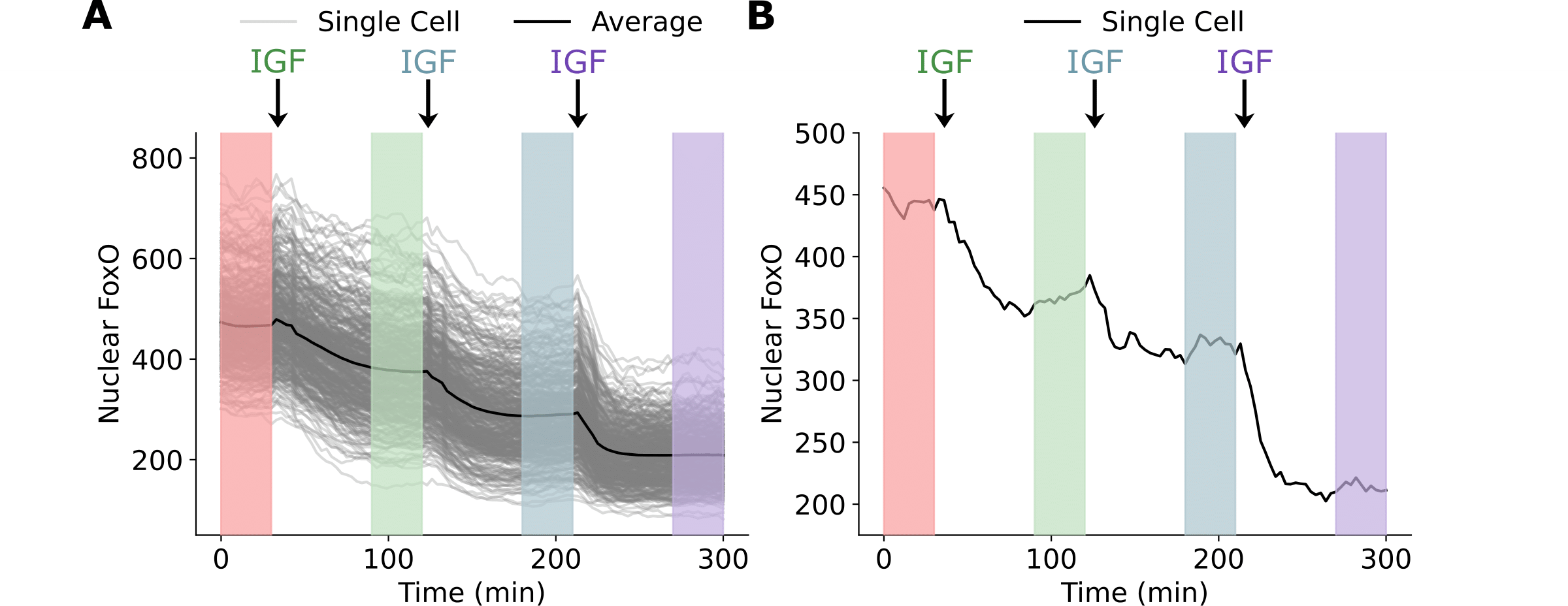

**Figure S6**. A schematic of the experiment performed by Gross et al. Single cells were treated with subsequent doses of IGF every 90 minutes and nuclear FoxO levels were recorded. FoxO levels reached a steady state at ~30-45 minutes of continuous IGF stimulation. Information about the steady state response was collected between 60 to 90 minutes of IGF stimulation (shaded regions). **A.** The single cell responses for all 400 cells examined in the step dose experiment along with the average response. **B.** The response of a single cell to the incremental dosing scheme.

Here, we describe how we estimated cell state specific mutual information from live cell imaging data. The IGF/FoxO pathway reaches an approximate steady state within 30-45 minutes after the introduction of the IGF ligand. We used data collected on HeLa cells where cells were treated with subsequent doses of IGF every 90 minutes to approximate the single cell response to multiple IGF levels. We approximated the single cell steady state nuclear FoxO distributions as gamma distributions and estimated their means and variances from data collected between 60 to 90 minutes of IGF stimulation (Fig. S6). Using these first two moments, we approximated the single cell response distribution $p(r|u,\boldsymbol{\theta})$ as a gamma distribution. The mutual information at the single cell level and the population average cell state specific mutual information $I_{Cee}$ were obtained using these response distributions. For $I_{CSA}$, the first two moments of the single cell response distributions were averaged to obtain the population level means and variances. This along with the assumption that the responses were gamma distributions provided the cell state agnostic response distribution $p(r|u)$ from which $I_{CSA}$ was obtained.

To verify that cell states are indeed conserved at the time scale of the experiment, we reanalyzed data generated by Gross et al.^9^ wherein cells were perturbed with IGF (37.5 pM), followed by a washout which allowed the cells to reach pre-stimulation nuclear FoxO levels, followed by a re-perturbation with the same amount of IGF. Nuclear FoxO response was measured at the single cell level after 90 minutes with IGF exposure both these times. Since the response $x$ to the same input $u$ was measured twice in the same cell ($x_{1}$ and $x_{2}$), we could evaluate the intrinsic variability in response at the single cell level. We then compared this intrinsic variability to the extrinsic cell-state dependent variability in the population.

To do so, we computed for each cell ${\delta=x}_{1}-x_{2}$ the difference between the two responses. SI Figure 7 show the histogram $p(\delta)$ as computed from the data (pink) and the same computed from the model that was trained on the single cell data (blue). We also computed ${p(\delta}_{0})$which represented the difference between responses of two different cells from the same population, shown for both data and the model.

As we see in SI Figure 7, the distribution $p(\delta)$ is significantly narrower than ${p(\delta}_{0})$ suggesting that intracellular variability is significantly smaller than across-population variability and that cells’ response to the same stimuli are quite conserved, especially when compared to responses in randomly picked pairs of cells. This shows that cell states and the corresponding response to extracellular perturbations are conserved, at least at the time scale of the experiment. Therefore, our estimates of cell-to-cell variability signaling fidelity are stable and reliable.

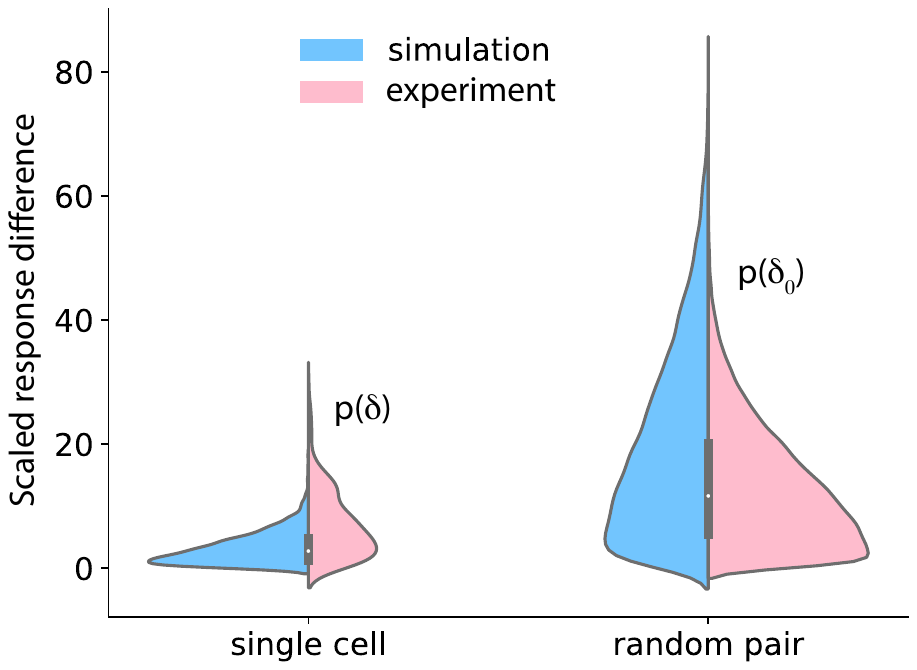

**Figure S7**. **Temporal stability of cell states.** Left: Cells were treated with 37.5 pM of IGF for 90 minutes, washed out for 120 minutes and again treated with 37.5 pM of IGF. Nuclear FoxO was measured during the treatment and the washout. The distributions on the left show the difference in FoxO levels in single cells after the two 90 minutes IGF stimulations (pink: data, blue: model). Right: Distribution of difference in FoxO levels in two randomly picked cells after 90 minutes of exposure to 37.5 pM IGF.

5. **Model predicted joint distributions** $p_{CeeMI}(I,\chi)$ **for several biochemical parameters**

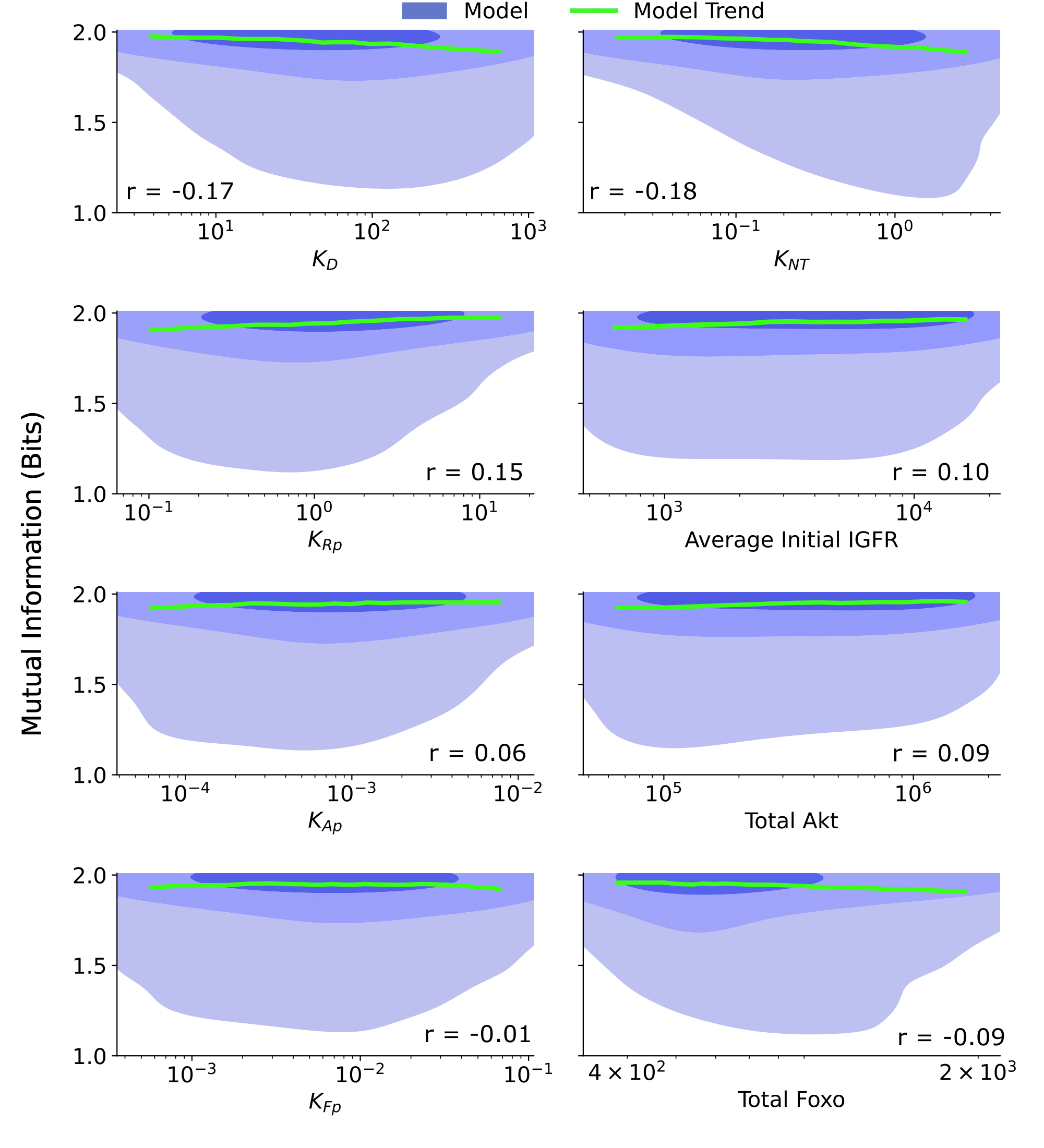

**Figure S8. Joint distributions for additional biochemical parameters.** The joint distribution $p_{CeeMI}\left( I,\chi\right)$ are provided for a collection of biochemical parameters: $k_{D}\equiv$ the ligand dissociation constant, $k_{NT}\equiv$ the ratio of the FoxO efflux rate to the FoxO influx rate, $K_{pI}\equiv$ the ratio of the IGFR phosphorylation rate to the pIGFR dephosphorylation rate, Average Initial IGFR $\equiv$ the average IGFR before IGF stimulation, given by the ratio of the production rate of IGFR to the degradation rate of IGFR, $K_{pA}\equiv$ the ratio of the Akt phosphorylation rate to the pAkt dephosphorylation rate, Total Akt $\equiv$ the combined count of pAkt and Akt, $K_{pF}\equiv$ the ratio of the cytoplasmic FoxO phosphorylation rate to the cytoplasmic pFoxO dephosphorylation rate, Total FoxO $\equiv$ the combined count of all forms of FoxO. The shaded blue regions are model predictions, and the green line is the model average. The correlation coefficient, r, is provided for each parameter. The contours represent 1% to 10%, 10% to 50%, and 50% to 100% of the total probability mass (from faint to dark shading).

**6. Examining the role of cell state dynamics on sensing ability**

To elucidate the role of cell state dynamics, we built a simple model of ligand-receptor system (main text Figure 4). We define the system as follows:

$$\begin{aligned} \phi\underset{\to}{k_{\mathrm{mprog}}}mRNA, mRNA\underset{\to}{k_{\mathrm{mdeg}}}\phi\#\left( S27 \right) \end{aligned}$$

$$\begin{aligned} mRNA\underset{\to}{k_{\mathrm{prod}}}mRNA+ R\#\left( S28 \right) \end{aligned}$$

$$\begin{aligned} R\underset{\to}{Lk_{\mathrm{bind}}}B, B\underset{\to}{k_{\mathrm{unbind}}}L+R\#\left( S29 \right) \end{aligned}$$

$$\begin{aligned} R\underset{\to}{k_{\deg}}\phi, B\underset{\to}{k_{\deg}}\phi\#\left( S30 \right) \end{aligned}$$

Here all cells are assigned identical kinetic parameters. Instead, the state of the system is defined by the abundances of each molecule prior to ligand dose. Thus, each cell may be described by two state variables, $\vec{\theta}=\{R_{0},mRNA_{0}\}$, where $mRNA_{0}$ is the initial mRNA levels and $R_{0}$ is the initial ligand-free receptor count. We tune the mRNA dynamics by changing mRNA production and degradation rates while keeping the average mRNA copy number constant. A relaxation of conditional responses requires both states have sufficiently high turnover, while we will modulate mRNA turnover, we will simply select a sufficiently high receptor turnover rate as described below.

For our simulations we set $\frac{kmprod}{kmdeg}=5 a.u.$, $k_{prod}=50 a.u. s^{-1}$, $k_{deg}=0.5 s^{-1},$and $k_{unbind}=1 s^{-1}$. We also set $k_{\mathrm{mdeg}}=\tau^{-1}$where $\tau$ takes on 5 values, equally distributed in the log space, between ${10}^{2}$ and ${10}^{4}$ s. We select two distinct cells to study, $\vec{\theta}_{a}=\{300,3\}$ and $\vec{\theta}_{b}=\{700,7\}$. For each cell and mRNA turnover timescale of interest, we introduce 20 different doses equally distributed in log scale leading to values of $Lk_{bind}$ ranging from ${10}^{-2}$ to ${10}^{3} (s^{-1})$. We ran 2,500 Gillespie simulations for each condition to obtain approximate distributions of the response. We then fit these samples to gamma distributions to obtain conditional responses from which the channel capacity mutual information can be straightforwardly calculated using an approach similar to the one described in SI Section 1.1.
